# Feeding behavior of *Aedes (Aedimorphus) vexans arabiensis*, a Rift valley fever virus vector in the Ferlo pastoral ecosystem of Senegal

**DOI:** 10.1101/593491

**Authors:** Biram Biteye, Assane Gueye Fall, Momar Talla Seck, Mamadou Ciss, Mariame Diop, Geoffrey Gimonneau

## Abstract

Host-vector contact is a key factor in vectorial capacity assessment and thus the transmission of mosquito-borne viruses such as Rift Valley Fever (RVF), an emerging zoonotic disease of interest in West Africa. The knowledge of the host-feeding patterns of vector species constitutes a key element in the assessment of their epidemiological importance in a given environment. The aim of this work was to identify the blood meal origins of *Aedes vexans arabiensis* Patton (*Diptera: Culicidae*), the main vector of RVF virus in the Ferlo pastoral ecosystem of Senegal. Engorged female mosquitoes were collected in Younouféré in the pastoral ecosystem in the Ferlo region during the 2014 rainy season. CDC light CO2-baited traps were set at six sites for two consecutive nights every month from July to November. Domestic animals present around traps were identified and counted for each trapping session. Blood meal sources of engorged mosquitoes were identified using a vertebrate-specific multiplexed primer set based on cytochrome b. Blood meal sources were successfully identified for 319 out of 416 blood-fed females (76.68%), of which 163 (51.1%) were single meals, 146 (45.77%) mixed meals from two different hosts and 10 (3.13%) mixed meals from three different hosts. *Aedes vexans arabiensis* fed preferentially on mammals especially on horse compared to other hosts (*P* < 0.001). Proportions of single and mixed meals showed significant temporal (*P* < 0.001) and spatial variations (*P* < 0.001) according to hosts availability. *Aedes vexans arabiensis* shows an opportunistic feeding behavior depending on the host availability. Results were discussed in relation with the Rift valley fever virus transmission and vector involvement as well as its primary hosts.

## Introduction

Rift Valley Fever (RVF) is an emerging zoonotic vector-borne viral infection [1] considered as a major problem of public and veterinary health as evidenced by various outbreaks in Africa [2–6]. This disease causes significant economic gaps in terms of animal deaths and economic losses in the affected countries [7–9]. Mosquitoes of the genera *Aedes* and *Culex* are the main vectors of RVF virus (RVFV) and transmission mainly occur during inter-epizootic periods [1]. RVF is endemic in Senegal, especially in the Ferlo region [10, 11]. The transmission of the virus is seasonal and insured by *Ae. vexans arabiensis* and *Cx. poicilipes* with peaks of transmission at the end of the rainy season [12–14]. Disease control is difficult because mosquito vectors are able to fly on long distances and escape the border sanitary barriers. Moreover, vector control methods are not used to control RVF outbreaks because they are costly and difficult to implement and could have important environmental and ecological consequences. However, host such as cattle could be treated with a remnant and efficient insecticide against the bites of mosquitoes, or parked at night in a fence surrounded by impregnated net might be used to reduce vectorial transmission in RVF outbreaks [15, 16].

The host-vector contact is a key factor in vectorial capacity assessment and the transmission of vector-borne pathogens. Understanding host-feeding pattern of vector species populations and their variations in space and time is important for a better knowledge of their respective roles in pathogens transmission, and thus in the design of accurate vector control measures or strategies [17]. Host choice is affected by innate preferences and environmental factors such as host diversity, density and distribution [18]. Although many studies on host preferences have been conducted for various mosquitoes, biting midges or tick vector species [17–22], so far in Senegal the molecular approach has been poorly used to determine the feeding behavior of disease vectors. Earlier investigations had used immunological assays [13, 19, 23] that have several inherent problems such as efficacy and reliability of blood meal identification [22, 24]. PCR based assays using different genetic markers have been developed for vectors targeting potential hosts for malaria, West Nile (WN) fever, African Horse Sickness or bluetongue research purposes (pig, human, goat, dog, cow; and avian) [17, 25–27]. PCR-based technology using host mitochondrial DNA provides a more direct approach to the identification of host species and increases sensitivity and specificity [22]. Mitochondrial DNA, particularly the cytochrome b (Cyt b), has been used extensively in various studies [28–31] because it exhibits a high level of interspecific polymorphism which helps to design species specific primers [32]. In this study, we have used a vertebrate-specific multiplexed primer set based on Cyt b to identify the blood meal origins of engorged females of *Ae. vexans arabiensis* caught during field collections. The aim of this work was to better understand the feeding behavior of RVFV vectors in the Ferlo pastoral ecosystem.

## Material and methods

### Study area

The study was performed around the Younouféré village (15°16’08.7”N and 14°27’52.5”W), a pastoral area located in the Ferlo region (central north of Senegal), during the 2014 rainy season. Younouféré is surrounded by small hamlets among which Diaby (15°17’18.1”N, 14°29’07.9”W), Demba Djidou (15°16’53.6”N, 14°27’04.8”W) and Nacara (15°13’23.1”N, 14°26’18.8”W) were selected as sampling sites (Fig 1). The area is characterized by a hot dry climate with a short rainy season (from June to October) and a long dry season (November to May), with mean annual rainfall ranging from 300 to 500 mm and a number of rainy days around 35.8 [33]. It is also characterized by a semi-arid steppe and many temporary ponds filled with rainfall and used by humans and animals as the main free sources of water during the rainy season [15, 34]. These ponds are the natural habitats of many species of birds, reptiles and rodents, and the breeding and resting sites for RVFV mosquito vectors. During the rainy season, the region became a high transhumance area where a high number of herds of domestic animals (cattle, sheep and goats) are concentrated around natural temporary ponds becoming thus risk areas due to the presence of vectors and the endemicity of the RVFV.

**Fig 1.**
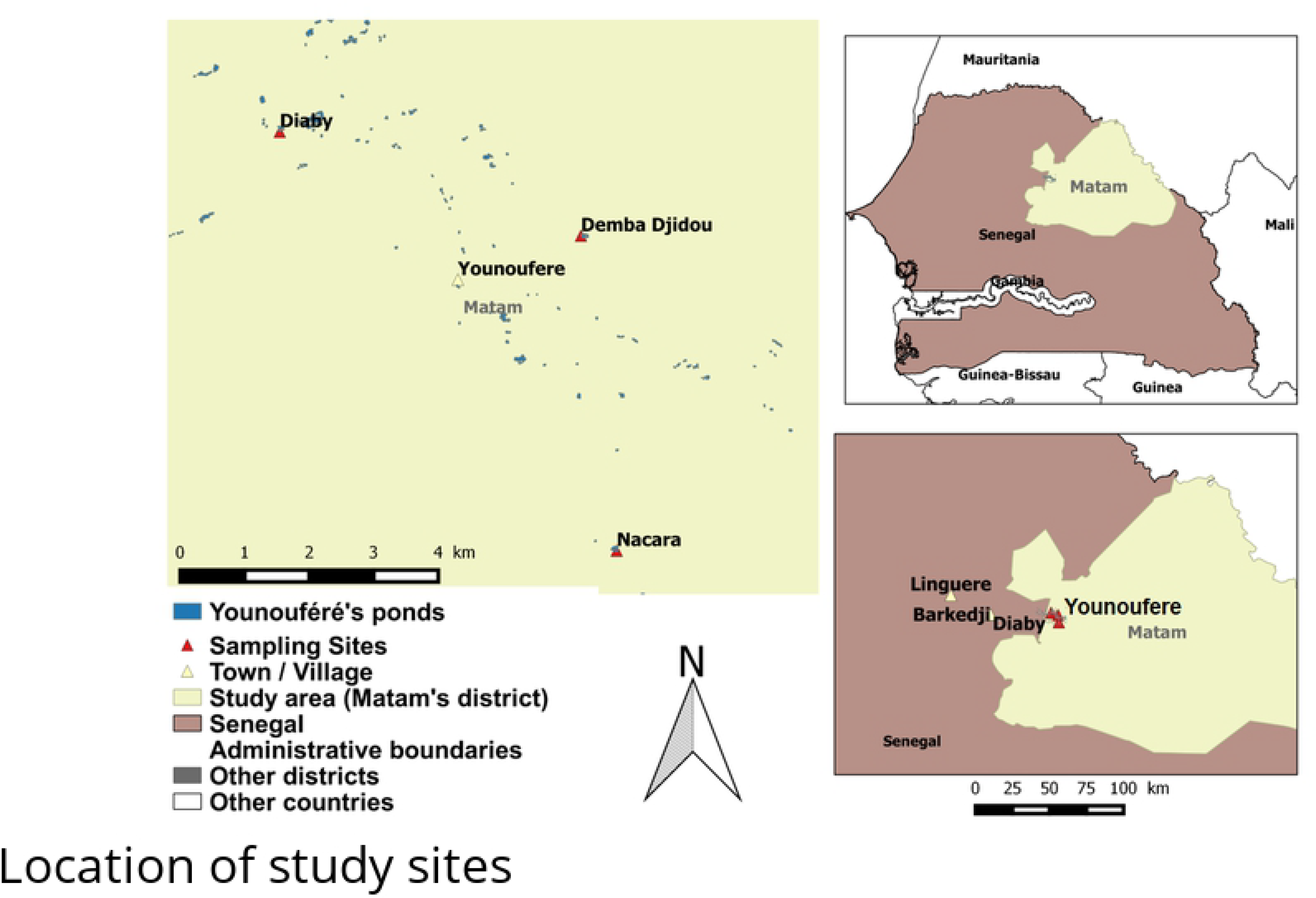
Location of the three sampling sites in Ferlo pastoral ecosystem (northern Senegal). Top-right corner: Senegal map and area of interest (in yellow). Bottom-right corner figure: triangles (in yellow) represent main towns/ villages nearby the sampling sites, while red triangles correspond to the sampling points. Main figure: detail of the positions of the three sampling points in Younouféré.

### Mosquito collection and survey of vertebrate hosts

Mosquito collection was conducted monthly during the 2014 rainy season from July to November. Mosquitoes were trapped nightly (from 6 PM to 6 AM) during two consecutive days in each study site, using miniature CDC light CO2-baited traps (BioQuip # 2836Q-6VDC, Rancho Dominguez, USA). Two traps were set per site at about 1.5m heights from the ground: one close to natural water point (ponds); another close to a herd pen. Distances between water source and herd pen varied from 100 to 800m depending on the site. In the field, the mosquitoes collected were killed by freezing in dry ice, sorted by genus on a chill table, put in 15 or 50 ml centrifuge tubes/cryo-tubes and transported in dry ice to the laboratory where they were identified according to sex and species on a chill table (−20 °C) using morphological keys [35, 36] and identification software [37–39]. The *Ae. vexans arabiensis* freshly engorged females were placed individually in eppendorf tubes (0.5ml) and stored at −20 °C until the analysis of the origin of the blood meals by PCR. Information on vertebrate hosts’ diversity and their relative abundance was recorded monthly around each trapping site (Fig 2). The presence of thousands of temporary pools also suggest a great diversity of wild fauna on which mosquito could feed. Thus the choice of primers used in this study was guided by the hosts identified around the sampling sites.

**Fig 2.**
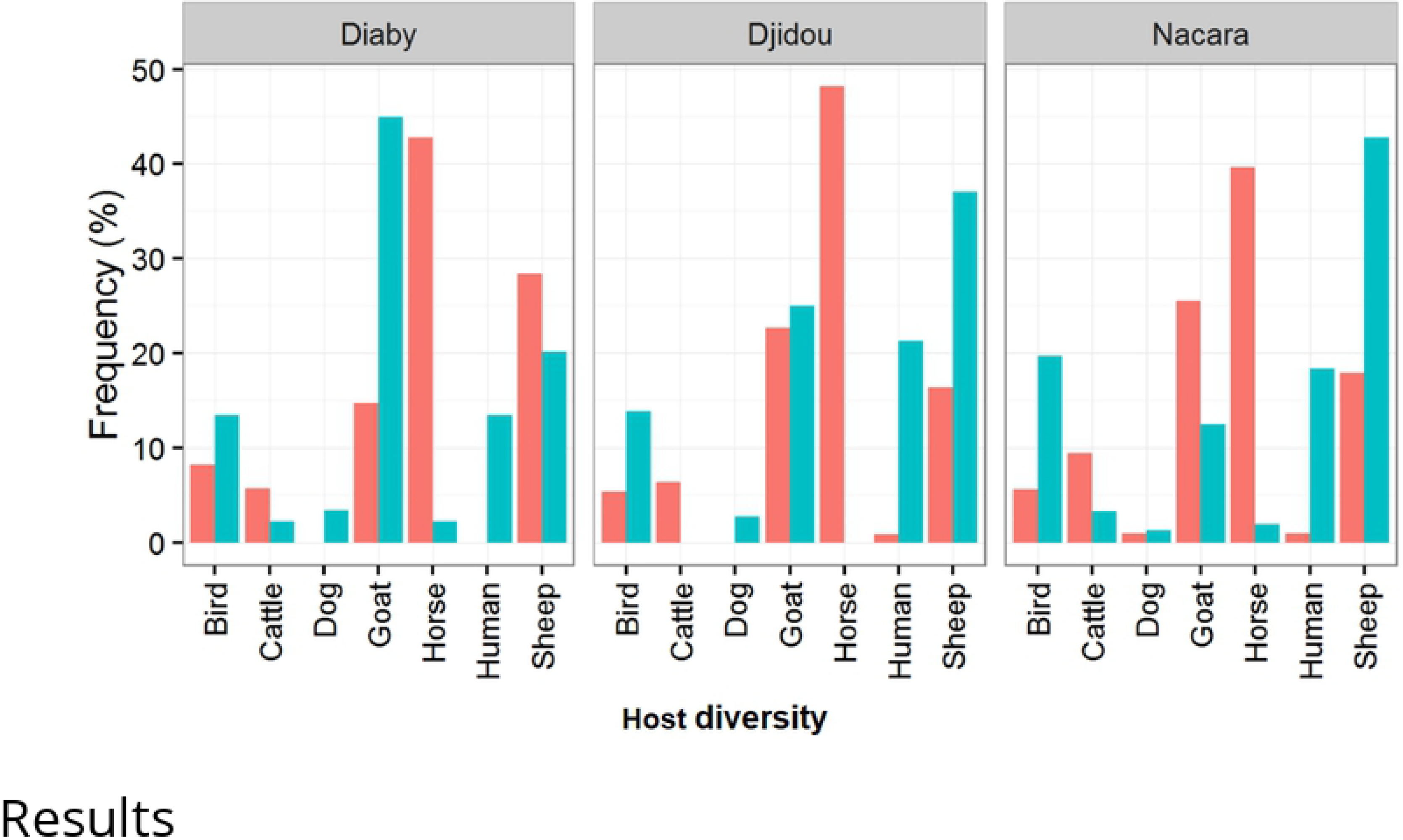
Origin of blood meals (%) taken by *Ae. vexans arabiensis* females collected at different sites (in pink) and vertebrate host proportions around trapping points (in cyan blue). On the x-axis we report the host diversity in each Site (column); on the y-axis the corresponding frequencies for each host.

### Extraction of genomic DNA and PCR amplification

Genomic DNA was individually extracted using the modified Chelex resin 10% extraction protocol (Resin Chelex100 ^®^, Chelating Ion Exchange Resin, Bio-Rad, France) described in [40–42]. Separated from the rest of the body, the mosquito’s abdomen was ground with sand using piston in an eppendorf tube of 1.5 ml. After grinding, a volume of 500 μl of Chelex solution was added to each tube. The tubes were incubated at 56 ° C for 2 hours (h) by vortexing every 30 minutes (min) and then at 95 ° C for 1 hour by vortexing every 20 min. Immediately after heating (thermal lysis), the tubes were centrifuged at 13,000 revs/min for 1 min to pellet the Chelex resin with inhibitor ions and cellular debris. The supernatants were gently transferred into new tubes and stored at −20 ° C until amplification of gene of interest.

Molecular identification was based on the amplification of the cytochrome b region of blood DNA as described in [17, 25, 26, 43, 44]. Two multiplex PCR were performed (Table 1) to separate cattle, sheep and goat first [17, 27] and dog and human on the other hand [43]. Two simplex PCR were used (Table 1) to identify blood meal from horse [17, 27] and from bird [26].

**Table 1.**
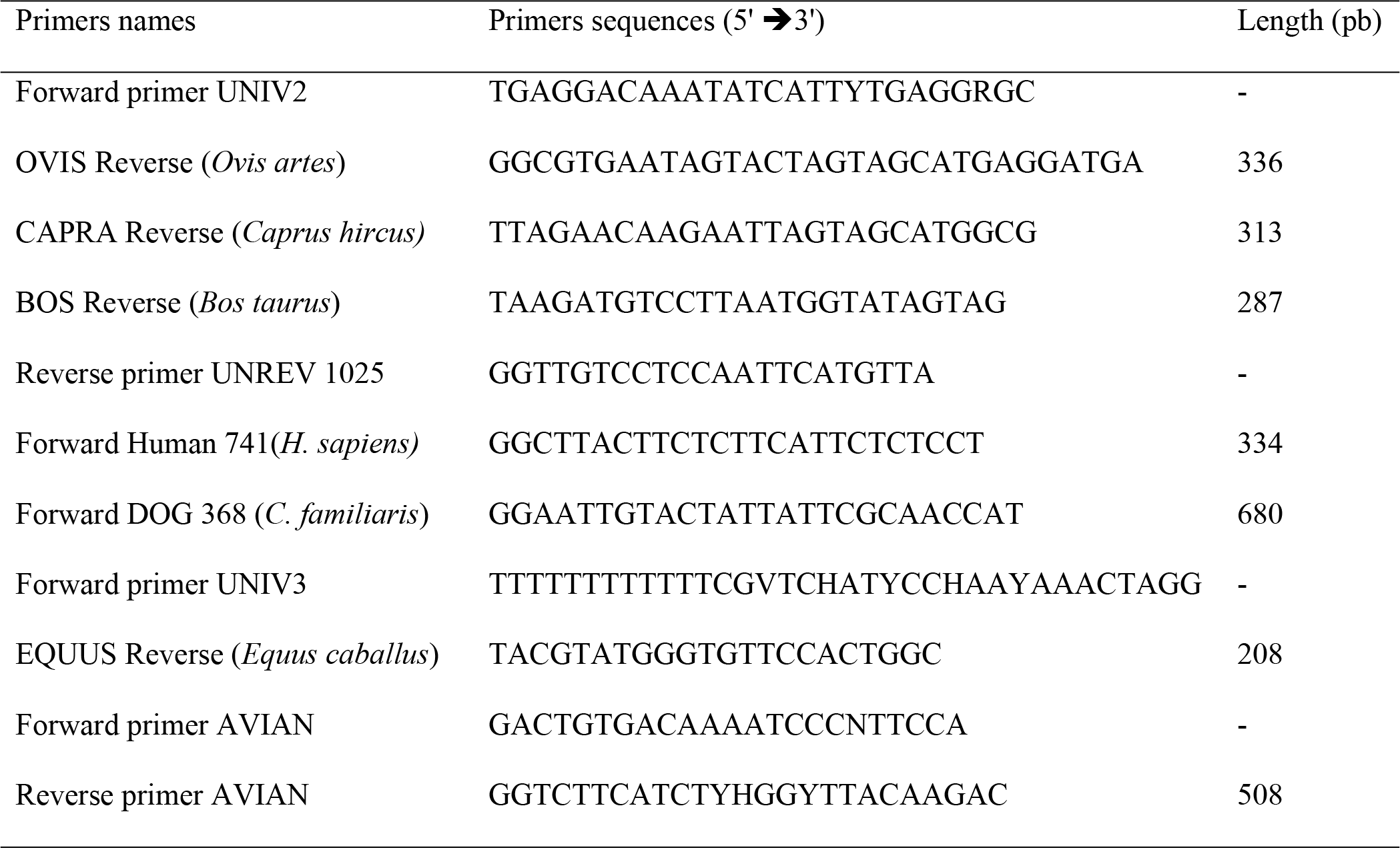
Primers set used for the identification of blood meal origin in mosquito abdomens.

### Statistical analysis

All night-traps allowed the collection of many specimens of mosquitoes of which a high number of engorged females of *Ae. vexans arabiensis*. However, a subsample was performed for the identification of blood meals origin given the high number of engorged females collected. Thus, for each month (from July to November) a maximum of 50 individuals was randomly selected from the data base for each trap point which collection exceeded 50 engorged individuals. And for each trap point which collection did not reach 50 engorged individuals, the total mosquito was analyzed. Non-parametric Kruskal-Wallis and Mann-Withney-Wilcoxon tests [45, 46] and the Chi-2 test (χ^2^) [47] were used to assess differences in engorged mosquitoes in time and between trap points. After descriptive analyses, a generalized linear model (GLM) was used to show whether there are preferential or opportunistic feeding behavior of *Ae. vexans arabiensis* depending on the host availability. All of the analyses were carried out using R software [48].

## Results

The 100 night-traps at 6 trap points allowed the collection of 104,352 specimens of *Ae. vexans arabiensis* of which 10,452 (10%) were engorged females. The subsampling carried out allowed the selection of 416 engorged females. Identification of blood meal sources was successful for 319 out of 416 blood-fed females (76.68%), of which 163 (51.1%) were identified as single meals, 146 (45.77%) as mixed meals from two different hosts and 10 (3.13%) as mixed meals from three different hosts. The 97 (23.31 %) remaining blood meals could not be determined with the primers used. According to single blood meals, *Ae. vexans arabiensis* fed preferentially on horse (n=73; 44.78%) and significantly (P < 0.001) more than goat (n = 45; 27.6%), sheep (n=22; 13.49%), cattle (n=10.42%) and bird (n=6; 3.68%); (Table 2; Fig 2).

**Table 2.** Poisson-GLM preference model used for *Ae. vexans arabiensis* vis-a-vis hosts.

|  | Regression coefficient | SE | Z-value | P-value |
| --- | --- | --- | --- | --- |
| Intercept | 4.26736 | 0.06836 | 62.426 | < 2e-16 |
| Bird | -1.81063 | 0.18233 | -9.930 | < 2e-16 |
| Cattle | -1.86947 | 0.18702 | -9.996 | < 2e-16 |
| Dog | -5.36598 | 1.00231 | -5.354 | 8.62e-08 |
| Goat | -0.83338 | 0.12420 | -6.710 | 1.95e-11 |
| Human | -4.67283 | 0.71040 | -6.578 | 4.78e-11 |
| Sheep | -0.61239 | 0.11530 | -5.311 | 1.09e-07 |

*Abbreviation:* SE, standard error

The number and type of meal at each trapping site are showed in Table 3. Of the 156 mixed meals, 141 (90.38%) involved horses; of which 131came from two different hosts and 10 coming from three different hosts. Of the 146 dual mixed meals, 81 (55.48%) concerned both sheep and horses, 36 (24.66%) concerned both goats and horses and only 3 (2.05%) concerned both sheep and goats. Only 2 mixed meals (1.28%) concerned humans showing a very high zoophilic rate. 6 single meals (3.68%) and 20 mixed meals (12.82 %) concerned birds which represents a high mammophilic rate.

**Table 3.** Origin of blood meals taken by *Aedes vexans arabiensis* females collected at different sites.

| Hosts | Diaby camp | Diaby Pond | Djidou Camp | Djidou Pond | Nacara Camp | Nacara Pond | Total |
| --- | --- | --- | --- | --- | --- | --- | --- |
| Goat | 12 | 7 | 3 | 9 | 5 | 9 | 45 |
| Cattle | 4 | 5 | 0 | 4 | 1 | 3 | 17 |
| Sheep | 9 | 4 | 1 | 1 | 3 | 4 | 22 |
| Human | 0 | 0 | 0 | 0 | 0 | 0 | 0 |
| Dog | 0 | 0 | 0 | 0 | 0 | 0 | 0 |
| Horse | 22 | 10 | 15 | 10 | 8 | 8 | 73 |
| Bird | 1 | 1 | 1 | 2 | 0 | 1 | 6 |
| Goat-Sheep | 0 | 1 | 0 | 1 | 0 | 1 | 3 |
| Goat-Cattle | 0 | 0 | 0 | 1 | 0 | 0 | 1 |
| Goat-Horse | 18 | 1 | 5 | 4 | 4 | 4 | 36 |
| Goat-Bird | 0 | 0 | 0 | 0 | 0 | 2 | 2 |
| Sheep-Horse | 53 | 4 | 13 | 1 | 6 | 4 | 81 |
| Sheep-Bird | 3 | 1 | 0 | 0 | 0 | 1 | 5 |
| Cattle-Horse | 3 | 1 | 0 | 1 | 1 | 4 | 10 |
| Cattle-Bird | 0 | 0 | 0 | 0 | 0 | 1 | 1 |
| Human-Horse | 0 | 0 | 1 | 0 | 1 | 0 | 2 |
| Horse-Bird | 0 | 1 | 1 | 0 | 0 | 0 | 2 |
| Goat-Sheep-Horse | 1 | 0 | 0 | 0 | 0 | 0 | 1 |
| Goat-Cattle-Horse | 0 | 0 | 0 | 1 | 0 | 0 | 1 |
| Goat-Dog-Horse | 0 | 0 | 0 | 0 | 0 | 1 | 1 |
| Goat-Horse-Bird | 1 | 0 | 0 | 0 | 0 | 1 | 2 |
| Sheep-Horse-Bird | 3 | 0 | 0 | 1 | 0 | 0 | 4 |
| Cattle-Horse-Bird | 1 | 0 | 0 | 0 | 0 | 0 | 1 |
| Total identified | 131 | 38 | 40 | 37 | 29 | 44 | 319 |
| Total unidentified | 27 | 17 | 17 | 12 | 18 | 6 | 97 |
| Total | 158 | 55 | 57 | 49 | 47 | 50 | 416 |

*Abbreviation:* Camp, livestock pen

The number of meals (all together) taken on the different hosts throughout the study (months) are presented in Table 4. The proportions of single blood meals and mixed blood meals (Fig 3) varied significantly over the time (χ^2^ = 258.1; *df* = 3, P < 0.001 and χ^2^ = 83.34; *df* = 3, P < 0.001 respectively); and between trapping points (χ^2^ = 21.9; *df* = 5, P < 0.001 and χ^2^ = 153.1; *df* = 5, P < 0.001 respectively). The highest abundances of single meals (n=129; 79.14%) and mixed meals (n=84; 53.85%) were observed in August.

**Table 4.** Variation of number of blood meals taken on hosts along the study period.

| Month | Total | Goat | Sheep | Cattle | Human | Dog | Horse | Bird | Mixed meals | Unidentified |
| --- | --- | --- | --- | --- | --- | --- | --- | --- | --- | --- |
| July | 7 | 2 |  | 0 | 0 | 0 | 0 | 0 | 5 | 0 |
| August | 294 | 37 | 13 | 13 | 0 | 0 | 60 | 6 | 84 | 81 |
| September | 56 | 3 | 4 | 3 | 0 | 0 | 7 | 0 | 31 | 8 |
| October | 59 | 3 | 5 | 1 | 0 | 0 | 6 | 0 | 36 | 8 |
| November | 0 | 0 | 0 | 0 | 0 | 0 | 0 | 0 | 0 | 0 |

## Discussion

The identification of the blood meals of hematophagous arthropods is very important in determining host-vector contact in nature. The PCR-based technology using host mitochondrial DNA has been used to eliminate some constraints of immunological assays [22] and to provide a more direct and sensitive approach to identify host species because sera do not have to be collected and specific antibodies produced [17]. In our study, the origin of the blood meal was successfully identified in 76.68% of the cases. Similar success rates have been observed in several studies [25, 49–51], while others get lower success rates [44, 52]. These different observations would probably be explained by the quantity and quality of the blood (partially digested or not), the diagnostic techniques, the range of primers used compared to the domesticated and/or wild fauna of the localities. Of the 319 meals identified, 156 (48.9%) were mixed meals and were mostly taken in August (53.85%). Similar observations have already been made by [19] and [23]. This high rate of mixed meals could be explained by the fact that August is the period corresponding to the peak abundance of *Ae. vexans arabiensis* [53] but also to the scarcity of hosts. In fact, this leads to the higher densities of vectors that are at the origin of a greater nuisance and consequently of a self-defense reflex in hosts. These self-defense reflexes developed by such few animals available and overexposed to the bites of a high population density of *Ae. vexans arabiensis*, fastly growing thanks to stocks of eggs from the previous rainy season, may even increase the rate of interrupted blood meals, favoring shifts between host species [23]. Thus, August seems to be the most favorable period for the spreading of the pathogens between hosts by *Ae. vexans arabiensis*.

Our results showed an opportunistic zoophagous behavior of *Ae. vexans arabiensis* populations that could feed on at least 7 different hosts. However *Ae. vexans arabiensis* showed a significant preference to feed on mammals and especially on horses (44.78%). In different proportions, this tendency to feed on horses has already been observed by [13, 19, 23] making this mosquito a potential bridge vector of the West Nile (WN) virus in the area [23]. Studies in Mexico and the northern United States (USA) also confirmed *Ae. vexans’* preference for mammalian vertebrates [51, 54]. Molaei & Andreadis [54] showed that populations of *Ae. vexans* mainly fed on deer (80%) and rarely on the horses (9.2%). It was mainly due to the fact that deer populations were more abundant and available than equine populations or simply by an acquired preference on deers. In our case, the large percentage of meals taken on horse is mainly explained by the fact that these animals are usually found at night in the immediate vicinity of the ponds where they find water and pasture and are the first available animals for active females for their blood meal. Therefore, the predominance of meals on horse could reflect a greater availability of the latter rather than a trophic preference compared to cattle and sheep. Although studies have shown that horses are not sensitive to RVFV infection and show a low viremia but the results suggest that they should be given particular attention for their high exposure to the virus transmission, as evidenced by significant prevalence (9.8%) in Nigeria [55] and the role they could play in the infection of vectors by co-feeding, a phenomenon already observed experimentally with the West Nile virus [56].

*Aedes vexans arabiensis* fed secondarily on small ruminants (goat and sheep), rarely on birds, humans and dogs, confirming the previous observations of [13, 19, 23] those of [57] and the important role that domestic ruminants probably play in the epidemiology of RVF. This observation would increase the risk of transmission of the RVFV, especially since the area is endemic for the disease. This would increase the vectors infection rate that can feed on susceptible small ruminants creating the occurrence or resurgence of the disease. Most authors showed a moderate or low rate of blood meal (0.4 to 10%) of *Ae. vexans arabiensis* on birds [23, 51, 54]. It should also be noted that most of the mixed meals (mainly recorded in August) originated from “horses-sheep” (55.48%) and “horses-goats” (24.66%). This finding is in line with the assumption that horses and domestic ruminants probably play a more important role in the epidemiology of RVF than the other hosts. The high rate of unidentified meals (23.32%) could be explained by the quantity and quality of the blood (partially digested or not), the presence in these localities of other groups of vertebrates such as rodents, reptiles, lagomorphs etc. that are present in the pools and that have not been tested in this study.

## Conclusion

The PCR-based technology using host mitochondrial DNA provides a more direct approach to the identification of host species and increases sensitivity and specificity compared to existing classical techniques. The blood meal sources of *Ae. vexans arabiensis* varied according to host availability and trap points and confirmed the opportunistic feeding behavior of the species. In the Ferlo pastoral ecosystem, horses and domestic ruminants probably play a more important role in the epidemiology of RVF than the other hosts. However complementary PCR tests, mainly focusing on wildlife of the mosquito breeding sites (pools) and sequencing need to be developed to elucidate their role in the maintenance of the RVF virus and to better understand the epidemiology of the disease in such localities where the disease is endemic.

## Acknowledgments

The authors are grateful to all the people who gave assistance in operating traps on several nights and surveys of hosts around the trapping sites. This study was partially funded by EU grant FP7-613996 Vmerge and is catalogued by the VMERGE Steering Committee as VmergeXXX (http://www.vmerge.eu). The contents of this publication are the sole responsibility of the authors and don’t necessarily reflect the views of the European Commission.

